# A Polymeric Nanoparticle System for the Delivery of CRISPR/Cas9 Components into Arabidopsis Pollen

**DOI:** 10.64898/2026.06.17.733037

**Authors:** Qinqin Yang, Liam D. Adair, Brian Jones, Markus Müllner

## Abstract

Efficient and heritable genome editing in plants remains constrained by transformation bottlenecks and reliance on tissue culture-based regeneration. Targeting the male germline offers a promising alternative for DNA-free genome modification. Here, we report a polymeric nanoparticle platform for the delivery of RNA-based CRISPR/Cas9 components into *Arabidopsis thaliana* pollen, establishing a foundation for sperm transfection-assisted genome editing (STAGE)-like approaches in plants. Poly(2-dimethylaminoethyl methacrylate) (PDMAEMA)-based polyplexes were designed to independently encapsulate Cas9 mRNA and ATTO 550-labeled guide RNA, forming nanoparticles with hydrodynamic diameters of ∼146 nm and condensed cores of 20-30 nm. Following internalization and cytosolic release, Cas9 mRNA translation enabled the nuclear localization of ATTO 550-labeled gRNA, as confirmed by confocal imaging and fluorescence lifetime (TauSense) analysis. Fluorescent signals corresponding to the CRISPR RNPs were detected in both vegetative and sperm cell nuclei, with higher accumulation in the vegetative nucleus. Together, these results demonstrate the feasibility of RNA-mediated delivery and intracellular assembly of CRISPR/Cas9 RNPs in plant male gametophytes. By bypassing tissue culture and DNA integration, this nanoparticle-based approach establishes a framework for a more efficient mechanism for introducing heritable genome modifications in plants.

**Graphical Abstract:** 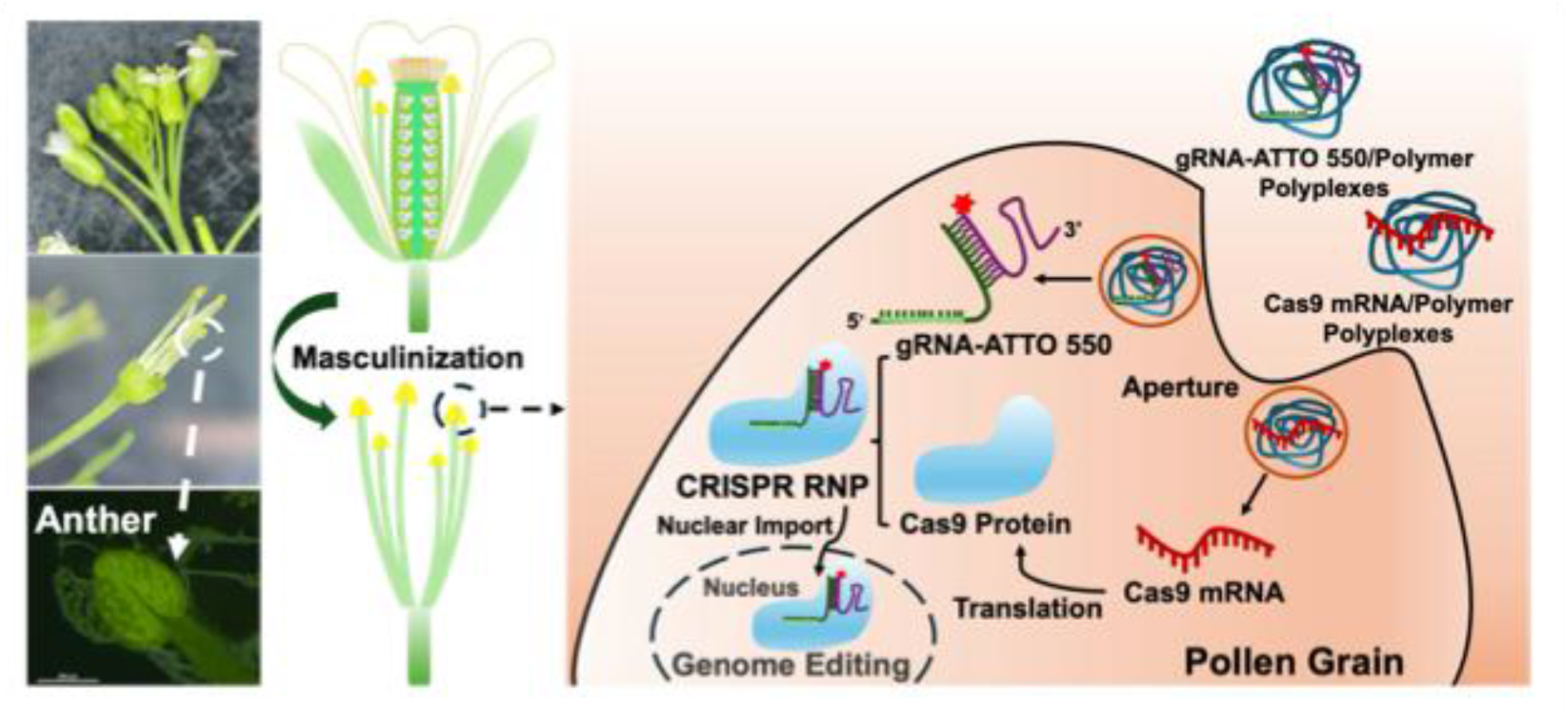

## Introduction

The continuous enhancement of productivity traits is a cornerstone of the modern agrifood system. CRISPR-based genome editing has begun to accelerate the process of trait improvement [1], but its application is limited by a co-dependence on the established plant transformation methods, *Agrobacterium*-mediated T-DNA delivery, protoplast transfection, and particle bombardment, that are labour-intensive and highly genotype-dependent [2]. In crop species, these approaches are largely coupled with tissue culture-based transformation of somatic cells, that further exacerbates genotype dependency and labour demands. Scaling CRISPR-based plant biotechnology requires new transformation methods that match the efficiency and broad efficacy of modern genome editing technologies.

Nanobiotechnology provides a promising approach to plant transformation, as it enables the design of nanomaterials that can carry functional cargo into cells. Polymer-based nanoparticles have the advantage of flexibility, enabling precise control over charge, size, and functionality. Cationic polymers, such as PDMAEMA, can form coacervate complexes with nucleic acids, condensing them into stable nanoscale carriers that can penetrate the plant cell wall matrix [3]. The development of an *Agrobacterium tumifaciens*-based method for targeting the female gamete has all but eliminated the need for somatic cell tissue culture-based plant transformation and subsequent whole plant regeneration in the model plant species, *Arabidopsis thaliana* [4]. The ability to transform female gametes via *in vivo* Agrobacterium infection is, however, largely limited to Arabidopsis. The egg cells in plants are deeply embedded in the complex pistil structure. By contrast, the male gametes in plants are contained within the pollen grains, that are generally abundant and readily isolated. Similarly to the sperm-mediated gene transfer (SMGT) and sperm transfection-assisted genome editing (STAGE) techniques in mammalian and other systems [5, 6], the male gametes in plants offer a compelling route for biomolecule delivery and heritable genetic modification. We have demonstrated that nanoscale coacervate PDMAEMA-based complexes can deliver functional nucleic acids into pollen cells, leading to the formation of CRISPR ribonucleoproteins (RNPs) that can localize to the pollen nuclei.

## Results & Discussion

### Polyplex Formation and Characterization

Previous studies have demonstrated nanoparticle-mediated delivery of biomolecules into pollen: including magnetofection using ∼200 nm poly(ethylene imine)-coated iron oxide nanoparticles for DNA [7]; or uptake of up to 50 nm layered double hydroxide nanoparticles for the delivery of short dsRNA (300 bp) [8]. We developed a system based on positively charged poly[2-(dimethylamino) ethyl methacrylate] (PDMAEMA) polymers for the delivery of biomacromolecules: EGFP mRNA, Cas9 mRNA, and ATTO 550-labeled gRNA. The PDMAEMA/biomacromolecule nanoscale coacervate complexes (i.e. polyplexes) were subsequently incubated with freshly harvested Arabidopsis pollen. PDMAEMA was selected due to its demonstrated nucleic acid binding capacity, pH-responsiveness, and suitability for intracellular delivery applications, as detailed in the Supplementary Information (**Figures S1-S3**). PDMAEMA contains pH-sensitive tertiary amine groups with an average pK_a_ of approximately 7.5, indicating that ∼50% of the amine groups are protonated and positively charged at physiological pH [9]. A pollen incubation media (PIM) was tailored to optimize both the extracellular stabilization of the polyplexes and pollen metabolic function. More details on the optimization of the PIM can be found in the Supplementary Information (**Figures S4 and S5**). At pH 5.8, PDMAEMA is strongly positively charged, while the Cap-1 modified Cas9 mRNA (Trilink) and ATTO 550-labeled gRNA (IDT) are negatively charged, facilitating coacervate formation (**Figure 1a**). A critical parameter determining the physicochemical properties of polyplexes is the ratio of the nitrogen atoms in the polymer (N) to the phosphate groups (P) in the nucleic acid backbone. Gel electrophoresis was used to monitor the formation of Cas9 mRNA and ATTO 550-labeled gRNA PDMAEMA polyplexes at different N/P ratios.

**Figure 1.**
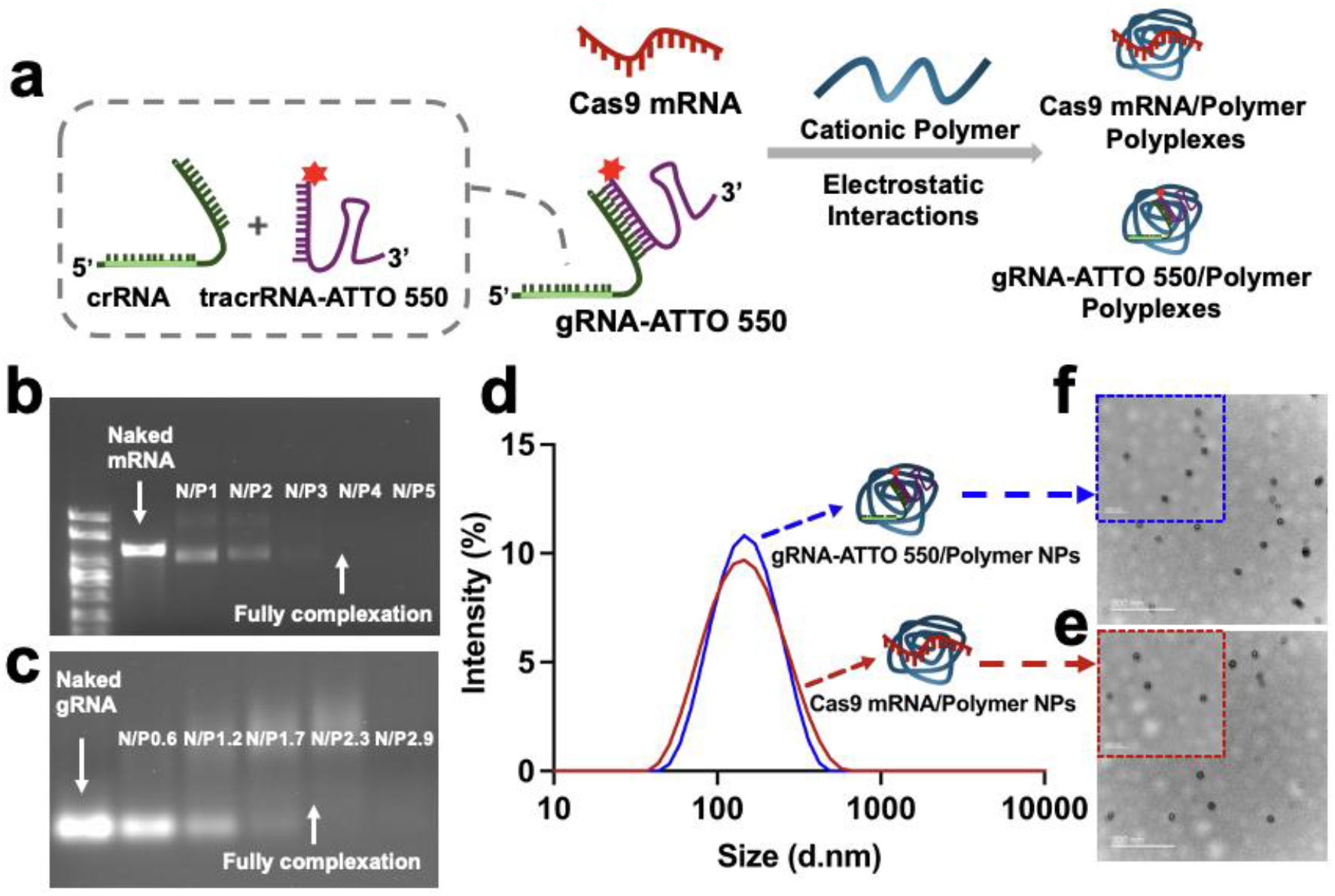
(**a**) Schematic illustration of polymer complexation with Cas9 mRNA or ATTO 550-labelled gRNA via electrostatic interactions. (**b**) Agarose gel electrophoresis analysis of Cas9 mRNA/polymer polyplex at different N/P ratios. (**c**) Agarose gel electrophoresis analysis of ATTO 550-labeled gRNA/polymer polyplex at different N/P ratios. (**d**) DLS measurements showing the intensity-based size distribution (D_H_) of Cas9 mRNA/polymer and ATTO 550-labeled gRNA/polymer polyplexes. (**e**) TEM image of Cas9 mRNA/polymer polyplex at N/P ratio 1. (**f**) TEM image of ATTO 550-labeled gRNA/polymer polyplex at N/P ratio 0.6. Scale bars represent 200 nm and 500 nm in (**e**) and (**f**).

For both Cas9 mRNA and ATTO 550-labeled gRNA, increasing the N/P ratio resulted in a progressive decrease in the intensity of free, non-complexed RNA bands, which ultimately disappeared, indicating enhanced polymer complexation with increasing N/P ratio. Notably, the Cas9 mRNA band was fully absent between N/P ratios of 3 and 4 (**Figure 1b**), while the gRNA band was completely undetectable between N/P ratios of 1.7 and 2.3 (**Figure 1c**), suggesting complete complexation within these ranges. As not all amine groups are fully protonated at physiological pH, increasing the N/P ratio provides a level to introduce more cationic charges within the system, promoting stronger interactions with negatively charged biomolecules (e.g. DNA, RNA and proteins).

The hydrodynamic diameter (D_H_) of the nanoscale complexes in aqueous suspension was determined by dynamic light scattering (DLS). At an N/P ratio of 1 for Cas9 mRNA polyplex and 0.6 for ATTO 550-labeled gRNA polyplex, well-defined size distributions were observed, with an average D_H_ of approximately 146 nm (**Figure 1d)**. The polyplex morphology was also characterized via transmission electron microscopy (TEM), indicating uniform and compacted nanostructures, where Cas9 mRNA (**Figure 1e**) and ATTO 550-labeled gRNA polyplexes (**Figure 1f**) each measured 20-30 nm.

### Internalization of Polyplex in Arabidopsis Pollen

The polyplexes can be internalized into pollen grains through the intine cell wall at apertures in the exine, and via endocytosis through the plasma membrane at these sites [10]. Cytoplasmic escape from endosomal vesicles is critical for the successful delivery of genetic cargo to the cytosol, nucleus and other subcellular compartments. Endosomal escape is typically triggered by pH changes within the endosome, which destabilizes the polyplexes, releasing the cargo into the cytosol (**Figure 2a**).

**Figure 2.**
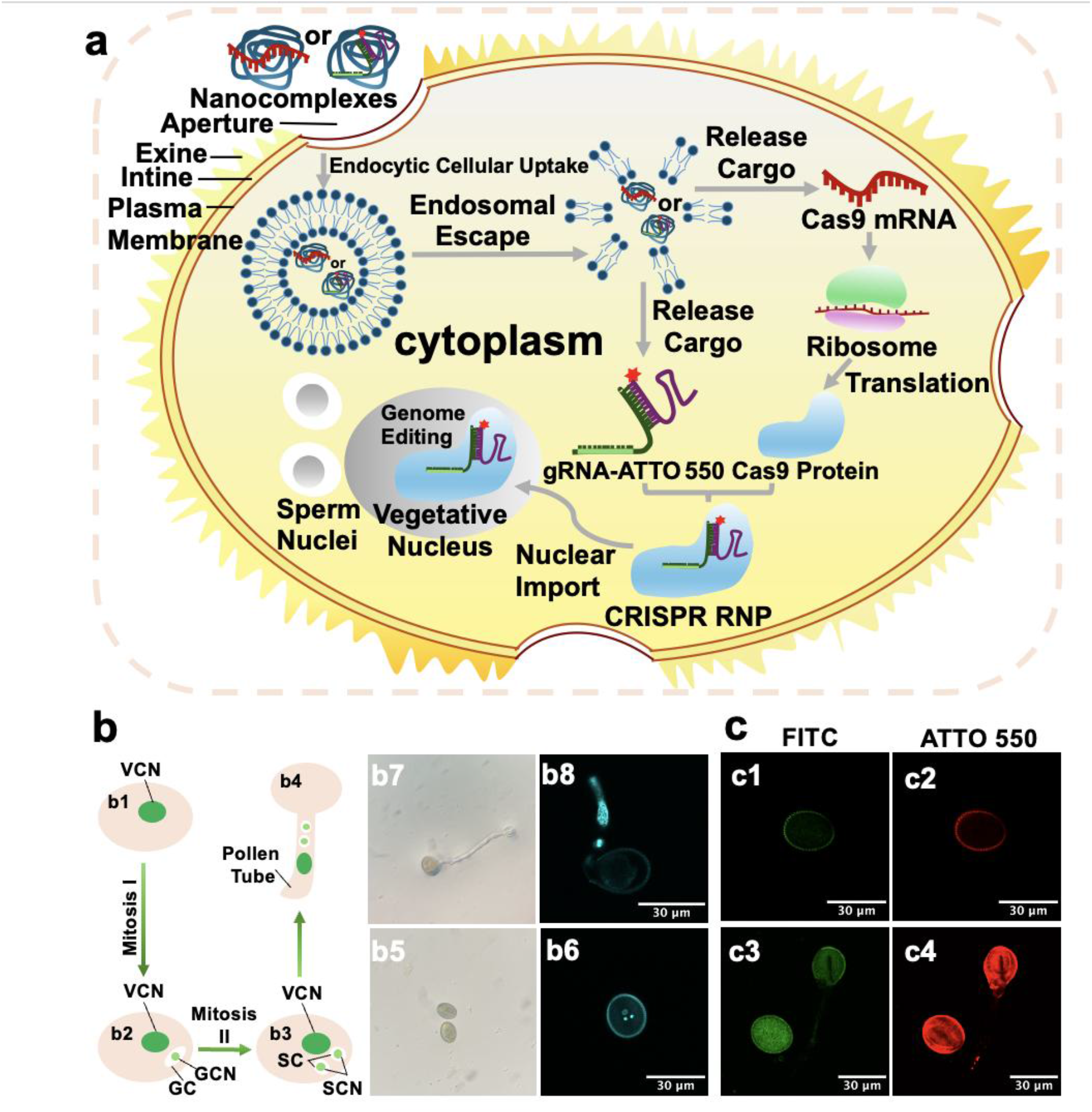
(**a**) Schematic overview of nanoparticle-mediated cargo delivery, intracellular cargo release, and subsequent functional activity. (**b**) Arabidopsis pollen at different developmental stages; (**b1**) Uninuclear pollen; (**b2**) Bicellular pollen; (**b3**) Trinuclear pollen with corresponding images in PIM (**b5**) and by confocal microscopy (**b6**), confocal image showing two sperm cell nuclei and a vegetative nucleus stained with Hoechst 33342 in the pollen grain; (**b4**) Germinated pollen with corresponding images in PIM (**b7**) and by confocal microscopy (**b8**), confocal image showing two sperm cell nuclei and a vegetative nucleus stained with Hoechst 33342 that have migrated into the pollen tube after germination. (**c**) Confocal microscopy images of pollen grains; (**c1-c2**) Untreated wild-type pollen; (**c3-c4**) Pollen treated with ATTO 550-labeled gRNA/Polymer polyplex. Scale bars represent 30 μm in (**b6**), (**b8**) and (**c1-c4**).

Each pollen grain in Arabidopsis develops three equidistant longitudinal apertures, which at the tetrad stage are aligned with apertures on sister microspores [11]. The developmental stages of the flower and anther are critical for successful pollen release and for ensuring that the pollen is mature. In Arabidopsis, mature pollen comprises the vegetative pollen cell and two fully internalised sperm cells are present at anther stage 12. Anther stages 10-12 correspond to floral developmental stages 11 and 12, during which pollen matures inside the locules. By anther stage 13, the stomium cells degenerate, allowing the mature pollen to be released. These later events align with flower stages 13 and 14, the stages used in this study [12]. The treatments consisted of a masculinisation of the male donor flowers, followed by nanoparticle treatments. Hoechst 33342 is a cell-permeable DNA stain that binds to A-T-rich regions in the minor groove of double-stranded DNA. It was used as a counterstain to enable visualization and identification of the sperm cell and vegetative nuclei (**Figure 2b**).

Freshly harvested, masculinized were first used to determine whether the relatively small ATTO 550-labeled gRNA cargo could be delivered into mature pollen via the polyplexes. Masculinized flowers were treated with ATTO 550-labeled gRNA polyplexes for 1 h and 20 min. Confocal microscopy of untreated pollen showed minimal autofluorescence (**Figures 2c1 and 2c2**), but a clear cytosolic ATTO 550 signal was detectable in the treated pollen (**Figures 2c3 and 2c4**), confirming successful transport of ATTO 550-labeled gRNA cargo across the hydrated pollen exine and intine cell walls and the plasma membrane. The delivery platform was then used to test for the ability to deliver the longer single-stranded, Cap-1 stabilized enhanced green fluorescent protein (EGFP) mRNA (Trilink) (**Figures S6-S8**). Confocal microscopy verified internal EGFP fluorescence in treated pollen, indicating successful polyplex delivery, and *in vivo* translation of the mRNA, and protein folding post treatment.

### *In Vivo* RNP Formation and Nuclear Co-localization

The Cas9 mRNA (Trilink) codes for a nuclear localized Cas9 protein. Several strategies were tested to determine if the uptake of the Cas9 mRNA polyplexes could result in the formation of Cas9/ATTO 550-labeled gRNA RNPs, as determined by the localization of ATTO 550 gRNA fluorescence in pollen nuclei. Hoechst 33342, a cell-permeable fluorescent stain, is widely used to label cell nuclei in live cells. The most efficient nuclear co-localization of ATTO 550-labeled gRNA with Hoechst 33342 (i.e. nuclear localization of CRISPR RNPs) was observed when the masculinized flowers were treated first with the Cas9 mRNA polyplexes, followed by an incubation with ATTO 550-labeled gRNA polyplexes (**Figure 3a**). The success of this sequential treatment aligns with previous findings where gene editing was enhanced when Cas9 mRNA was introduced first, allowing sufficient time to produce the Cas9 protein before the cellular delivery of gRNA. The ability to capture gRNAs in stable RNPs complexes presumably reduces premature gRNA degradation [13].

**Figure 3.**
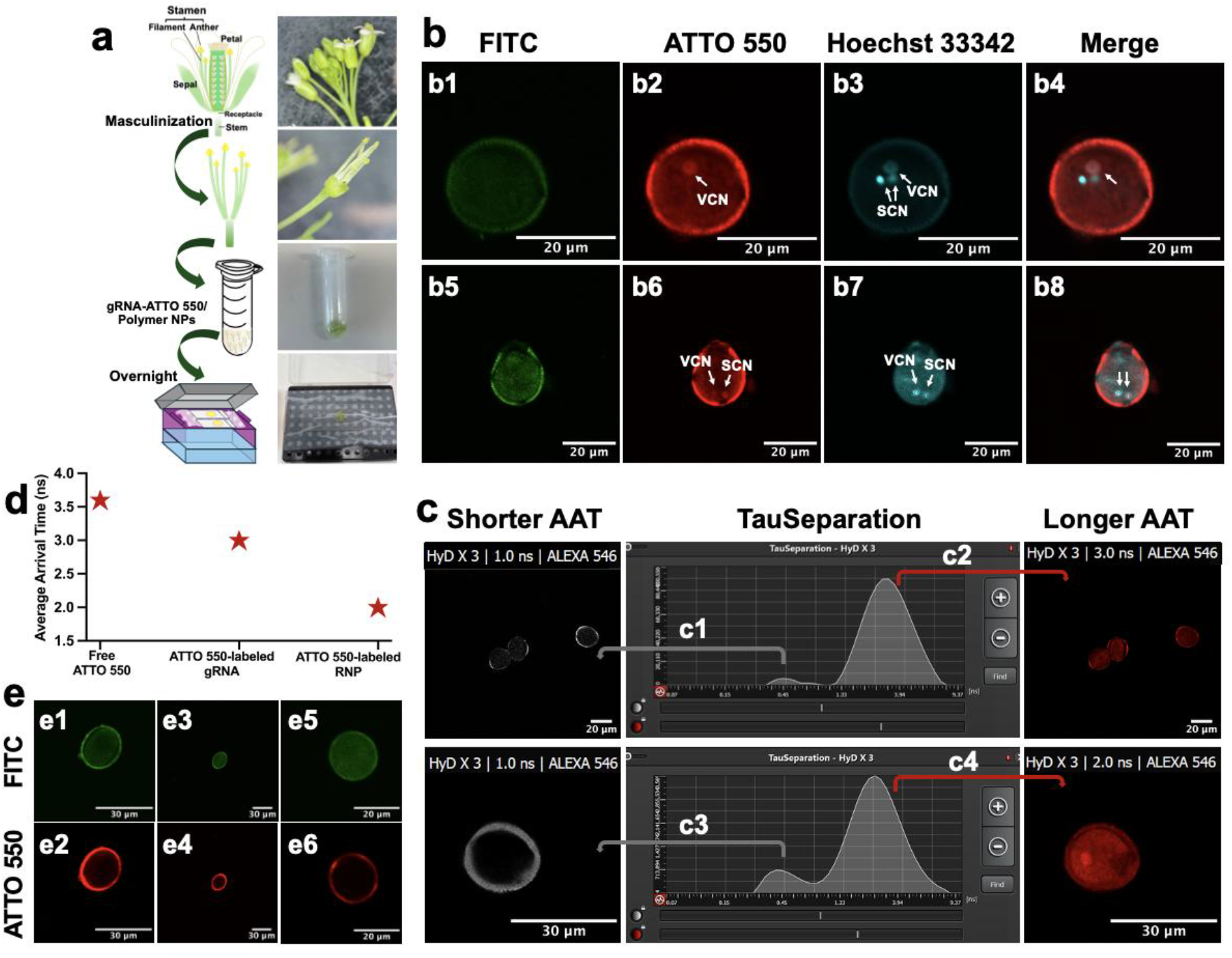
(**a**) Schematic illustration of the treatment schemes and sequential application of Cas9 mRNA/polymer polyplex and ATTO 550-labeled gRNA/polymer polyplex. (**b**) Confocal images of pollen treated with Cas9 mRNA/polymer polyplex (N/P ratio 1) followed by overnight incubation with ATTO 550-labeled gRNA/polymer polyplex (N/P ratio 0.6); ATTO-550 signal is shown co-localized with Hoechst 33342-stained nuclei. (**c**) Tau separation analysis of ATTO 550 signal in Arabidopsis pollen; (**c1-c2**) ATTO 550-labeled gRNA signal; (**c3-c4**) Signal corresponding to Cas9 protein/ATTO 550-labeled gRNA RNP formation. (**d**) Comparison of average arrival time of free ATTO 550, ATTO 550-labeled gRNA and ATTO 550-labeled RNP. (**e**) Confocal images of different polyplex treatment conditions; (**e1-e2**)Pollen treated with ATTO 550-labeled gRNA/polymer polyplex followed by overnight incubation with Cas9 mRNA/polymer polyplex; (**e3-e4**) Pollen co-treated with ATTO 550-labeled gRNA/polymer and Cas9 mRNA/polymer polyplexes in PIM overnight; (**e5-e6**) Pollen treated with combined ATTO 550-labeled gRNA and Cas9 mRNA/polymer polyplexes in PIM overnight. Scale bars represent 20 μm and 30 μm in (**b**), (**c**) and (**e**).

Hoechst 33342 counterstaining typically enables the visualization of all three nuclei in pollen grains, the two sperm cell nuclei and the vegetative nucleus. The formation of RNPs complexes with the ATTO 550-labeled gRNA, localized predominantly to the vegetative nucleus and occasionally additionally to sperm cell nuclei. In most pollen grains, the ATTO 550 signal was detected in the vegetative nucleus, with no detectable co-localization in sperm nuclei (**Figure 3b4**). In a subset of pollen grains, however, the ATTO 550 signal was observed in both the vegetative nucleus and sperm nuclei, overlapping with Hoechst 33342 staining (**Figure 3b8**).

Confocal microscopy combined with tau-based fluorescence lifetime analysis further confirmed the presence of the ATTO 550 signal within the vegetative nucleus. Pollen exhibits intrinsic autofluorescence with an average arrival time (AAT) of approximately ∼1.0 ns [14], whereas free ATTO 550 shows an AAT of ∼3.6 ns. Tau separation analysis was performed on pollen samples treated with (i) ATTO 550-labeled gRNA polyplex and (ii) Cas9 mRNA polyplex followed by incubation with ATTO 550-labeled gRNA polyplex. In both cases, two distinct AAT populations were resolved. A short component (∼1.0 ns), corresponding to endogenous pollen autofluorescence, was consistently observed (**Figure 3c1 and 3c3**). A longer component corresponding to ATTO 550 signal showed an AAT of ∼3.0 ns for ATTO 550-labeled gRNA released from polyplex in the cytoplasm (**Figure 3c2**), and ∼2.0 ns for Cas9 protein/ATTO 550-labeled gRNA RNP (**Figure 3c4**). Overall, the AAT decreased progressively from 3.6 ns (free ATTO 550) to 3.0 ns (gRNA-associated ATTO 550) and further to 2.0 ns upon RNP formation (**Figure 3d**). This systematic shift likely reflects changes in the local microenvironment of the fluorophore upon protein binding and intracellular delivery into the nucleus. Protein association and the surrounding nuclear environment may influence fluorescence properties through effects such as altered solvent accessibility, changes in local polarity, and potential quenching interactions, which can collectively contribute to reduced fluorescence lifetime compared with the free dye in solution. These results further confirm the successful nuclear delivery of *de novo* synthesised RNPs.

In contrast to the results of the sequential delivery of Cas9 mRNA/polymer polyplex followed by ATTO 550-labeled gRNA/polymer polyplex, other delivery strategies did not produce noticeable nuclear co-localization (**Figure 3e**). Successful nuclear co-localization of the ATTO 550-labeled gRNA with Hoechst-stained nuclei was only observed when stamens were first incubated with Cas9 mRNA/polymer polyplex for 1 h 20 min, followed by overnight incubation with gRNA-ATTO 550/polymer polyplex. This sequential delivery strategy likely allowed sufficient time for internalized Cas9 mRNA to escape the endosomal compartment and undergo cytosolic translation prior to the introduction of gRNA-ATTO 550. The resulting Cas9 protein, containing nuclear localization signals would then be available to assemble with the subsequently delivered ATTO 550-labeled gRNA to form functional RNP complexes capable of nuclear import. When ATTO 550-labeled gRNA/polymer polyplex was delivered before Cas9 mRNA/polymer polyplex, the ATTO 550 signal was primarily detected at the pollen exine and did not appear within the pollen cytoplasm observed (**Figure 3e1 and 3e2**). A comparable pattern was observed when pollen grains were simultaneously exposed to ATTO 550-labeled gRNA/polymer polyplex and Cas9 mRNA/polymer polylpex in PIM observed (**Figure 3e3 and 3e4**). In these treatments, the ATTO 550 signal accumulated on the pollen surface, suggesting that ATTO 550-labeled gRNA delivery was impeded under these conditions. The absence of internal fluorescence may be related to changes in nanoparticle stability or aggregation behaviour in the PIM, that contains ions and other components known to influence nanocomplex surface charge and cellular uptake efficiency. Furthermore, co-delivery of Cas9 mRNA and ATTO 550-labeled gRNA within the same polymeric nanocomplex did not produce detectable ATTO 550 signal either on the exine or within the pollen grains observed (**Figure 3e5 and 3e6**). This observation may reflect altered complexation behaviour when two nucleic acid cargos compete for interaction with the cationic polymer, potentially leading to unstable or oversized complexes that are less efficiently internalized. Previous studies of cationic polymer-mediated nucleic acid delivery have shown that cargo composition, charge ratio, and complex architecture strongly influence nanoparticle size, stability, and uptake efficiency [15, 16]. Taken together, these results highlight the importance of temporal control of cargo delivery for successful intracellular assembly of CRISPR/Cas9 components in pollen. The sequential delivery strategy employed here enables intracellular production of Cas9 protein prior to ATTO 550-labeled gRNA uptake, thereby facilitating efficient RNPs formation and nuclear import. These findings suggest that optimizing the timing and composition of polymer-nucleic acid nanocomplexes is a critical parameter for achieving functional delivery of genome editing reagents into plant reproductive cells.

## Conclusions

In summary, this report describes the development of a polymer-based nanoparticle platform for delivering functional RNA into *Arabidopsis thaliana* pollen. The predominance of localization in the vegetative nucleus indicates that nuclear import is highly efficient in the vegetative cell. The reduced frequency of localization to the sperm cell nuclei highlights a potential for further refinement of the nuclear localization signal sequences incorporated into the Cas9 mRNA and protein. The results reveal that once released from the polyplexes in the pollen cytosol, Cas9 mRNA can be translated and form nuclear localized Cas9 protein/ATTO 550-labeled gRNA RNPs. Delivery to sperm cell nuclei establishes a potential foundation for male gamete-driven precision genome editing in plants, either directly via the gametes or via delivery to the zygote in fertilization, laying the groundwork for optimizing heritable plant genetic modification and direct, tissue-culture-free route genome editing in plants.

### Experimental Section Materials

Chemical reagents used in the studies consisted of: CleanCap^®^ Cas9 mRNA and CleanCap^®^ EGFP mRNA (Trilink), CRISPR-Cas9 crRNA and CRSPR-Cas9 tracrRNA-ATTO 550 (IDT), agarose gel (Bioline), SYBR™ Safe DNA Gel Stain (Invitrogen), λ DNA ladder (Thermo Fisher Scientific), RNA Markers (Promega), microRNA Markers and 2× RNA Loading Dye (New England Biolabs) were purchased for the study. Sucrose (Ultrapure; Biochemicals), boric acid (Astral Scientific), calcium nitrate (ChemSupply), potassium chloride (Ajax), magnesium sulphate (Mallinckrodt), ethylenediaminetetraacetic acid (EDTA; Merck), acetic acid (Biolab), hydroxymethyl methylamine (Tris; ChemSupply).

## Methods

### *Arabidopsis Thaliana* Growth Conditions

The WT *Arabidopsis thaliana* ecotype Columbia-0 (Col-0) was used in this research. Col-0 seeds were obtained from the ABRC [17]. Plants were grown in a growth room maintained at a constant 22.5 °C under an 18 h light/6 h dark photoperiod.

### *In Vitro* Pollen Germination Test

The *in vitro* PIM was optimized to support Arabidopsis pollen metabolism, with the following composition: 15% (w/v) sucrose, 1.6 mM boric acid, 1.7 mM calcium nitrate, 1.0 mM potassium chloride, and 1.0 mM magnesium sulphate, adjusted to pH 5.8 using 0.1 M KOH or HCl. To assess the potential cytotoxic effects of the nanocomplexes on pollen viability, *in vitro* pollen germination assays were conducted using an improvised humidity chamber. This chamber consisted of an empty pipette tip box lined with a moist Kimwipe and a shallow layer of water to maintain humidity. A 30 μL drop of PIM was placed on a glass slide, and pollen grains were carefully transferred into the medium. The prepared slides were incubated in the humidity chamber in the dark overnight at room temperature. Pollen germination was evaluated using a Leica DM2700 M light microscope.

### Complexation of crRNA and tracrRNA

crRNA and tracrRNA-ATTO 550 (IDT) oligonucleotides were each resuspended to a stock concentration of 100 µM in nuclease-free duplex buffer. To generate gRNA, crRNA and tracrRNA-ATTO 550 were mixed at a 1:1 molar ratio to obtain a final duplex concentration of 100 µM. The mixture was heated at 95 °C for 5 min and then allowed to cool to room temperature (∼15 min) to facilitate duplex formation. The resulting crRNA:tracrRNA gRNA duplexes were stored at −30 °C until further use.

### Conjugation of Cas9 mRNA and ATTO 550-labeled gRNA with Polymer

To prepare the nanocomplexes, predetermined amounts of PDMAEMA were added to pH 5.8 PIM buffer in 1.5 mL microcentrifuge tubes, followed by the dropwise addition of a fixed amount of Cas9 mRNA. A molecular mass of 330 Da per mRNA phosphate group was assumed in N/P ratio calculations. The mixtures were vortexed vigorously for 10 s and incubated at room temperature in the dark for 30 min. The resulting polyplexes were characterized using 1% agarose gel electrophoresis, DLS, and TEM. To prevent nanoparticle aggregation, fresh nanocomplex solutions should be prepared immediately prior to use in pollen handling experiments. The ATTO 550-labeled gRNA was conjugated to the polymer using the same protocol as described for the Cas9 mRNA conjugation above.

### Agarose Gel Electrophoresis

All DNA fragments were visualized using a 1% (w/v) agarose gel prepared in 1× TAE buffer and stained with SYBR™ Safe DNA Gel Stain. Samples were mixed with 2X RNA Loading Dye prior to loading. RNA markers were similarly prepared with 2× RNA loading dye, heated at 67 °C for 10 min, and immediately cooled on ice for 2 min before loading. microRNA was heated at 67 °C for 10 min and immediately cooled on ice for 2 min before loading. Electrophoresis was performed at 80 V for 60 min. Gels were subsequently visualized using a UV transilluminator.

### Dynamic Light Scattering

DLS was used to determine the hydrodynamic diameter and polydispersity index of the polyplexes. Measurements were performed using a Zetasizer Nano Series (Malvern Instruments). All samples were syringe-filtered. Freshly prepared samples were equilibrated at 25°C for 2 min to optimize the measurements.

### Transmission Electron Microscopy

The morphology and particle size distribution of the polyplexes were analysed using TEM on a FEI Tecnai F12 operating at 120 kV in brightfield mode. Prior to sample loading, TEM grids (300 mesh, carbon-coated) were rendered hydrophilic through a 30-second glow discharge using a Dina Plasma Technology system, enhancing sample adsorption. Freshly prepared polyplex solutions (3 μL) were applied onto the treated grids and allowed to adsorb for 1 min. Excess liquid was gently blotted with filter paper, and the grids were washed three times with Milli-Q water. Samples were then negatively stained twice using 3 μL of 2% uranyl acetate for 45 s each. After staining, the grids were air-dried at room temperature for 1 minute before imaging. TEM images were processed and analysed using ImageJ software, applying grayscale analysis to assess particle morphology and size distribution.

### Pollen Grain Nanocomplex Uptake and Treatment

Cas9 mRNA/polymer nanocomplex solution was prepared at an N/P molar ratio of 1. Flowers at developmental stages 13-14 were emasculated under a stereomicroscope using fine tweezers, and thirty flowers were transferred into sterile 2.0 mL microcentrifuge tubes. Cas9 mRNA/polymer nanocomplexes were added to fully immerse the floral tissues. Samples were briefly centrifuged at 4200 rpm for 10 s to remove trapped air bubbles and incubated at room temperature on a shaker (100 rpm) for 1 h 20 min in the dark.

Following incubation, stamens were washed twice with 200 µL Milli-Q water per wash to remove excess nanoparticles. Nuclear staining was performed by adding Hoechst 33342 to a final concentration of 0.8 mg/mL. Samples were again centrifuged briefly to remove air bubbles and incubated at room temperature on a 100-rpm shaker for 1 h in the dark.

After Hoechst staining, stamens were washed twice with 200 µL Milli-Q water to remove excess stain and gently blotted dry with Kimwipes. For ATTO 550-labeled gRNA delivery, 100 µL of ATTO 550-labeled gRNA/polymer nanocomplex (N/P ratio 0.6) solution was added to a well microscope slide, and the stamens were placed into the well. An additional 100 µL of nanocomplex solution was added to ensure complete coverage. Slides were then transferred to a humid chamber constructed from a pipette tip box containing a shallow layer of water and lined with a moist Kimwipe to maintain humidity. The chamber was covered with foil, and samples were incubated overnight (∼22 h) at room temperature in the dark to allow transgene expression.

The following day, stamens were washed twice with 200 µL Milli-Q water directly in the well to remove residual nanoparticles and gently blotted dry. Pollen grains were collected by tapping the stamens onto microscope slides pre-coated with PIM, and coverslips were placed on top for imaging.

Confocal imaging was performed using an Olympus FV3000 laser scanning confocal microscope with a 40× objective or a Leica Stellaris 8 confocal and multiphoton microscope with a 25× water-immersion objective. Z-stack imaging was used to acquire optical sections through pollen grains and assess whether fluorescence signals were localized within or outside the cells. Image analysis was conducted using Fiji software (version 2.14.0/1.54f).

## Author Contributions

M.M. and B.J. jointly conceived the study and supervised the work. Q.Y. planned and developed the experimental setup, performed all experiments, and analyzed the data. L.D.A. synthesized the clickable dye. Q.Y. drafted the manuscript with input from all co-authors.

## Acknowledgments

This research was facilitated by access to Sydney Analytical, a core research facility at the University of Sydney. The authors acknowledge the technical and scientific assistance from Sydney Microscopy & Microanalysis (SMM). The authors thank Key Centre for Polymers & Colloids (KCPC) for access to equipment. Q.Y. is a grateful recipient of Henry Bertie and Florence Mabel Gritton Research Scholarship from the University of Sydney.

## Data Availability Statement

Data is provided within the manuscript and supplementary information files.

